# Coexisting with humans: genomic and behavioural consequences in a small and isolated bear population

**DOI:** 10.1101/2025.05.15.654188

**Authors:** Giulia Fabbri, Roberto Biello, Maëva Gabrielli, Sibelle Torres Vilaça, Beatrice Sammarco, Silvia Fuselli, Patrícia Santos, Lorena Ancona, Laura Peretto, Giada Padovani, Marco Sollitto, Alessio Ianucci, Ladislav Paule, Dario Balestra, Marco Gerdol, Claudio Ciofi, Paolo Ciucci, Carolyn G. Mahan, Emiliano Trucchi, Andrea Benazzo, Giorgio Bertorelle

## Abstract

Climate and land use change have increased human-wildlife interactions, potentially reducing wild species density and prompting behavioural adaptations to urbanised environments. It is still debated if behavioural responses are mainly the result of phenotypic plasticity or if they were driven by anthropic selective pressures, especially in small populations. Our study focused on the Apennine brown bear population (*Ursus arctos marsicanus*), which has coexisted with humans in Central Italy for millennia. We characterised genomic diversity and identified adaptation signals distinctive to this population by comparing whole genome resequencing data across the Holarctic species range. We show that Apennine brown bears possess a unique genomic diversity pattern including selective signatures at genes associated with reduced aggressiveness, possibly involving alternative splicing mechanism. Our findings suggest that even in small and long-isolated populations, selection may shape behavioural traits. We hypothesise that human-induced selection has influenced these changes, reducing conflicts and contributing to the long-term persistence of the Apennine bear and its coexistence with humans.

## MAIN

Humans have long influenced the environment in which they lived and thrive, affecting ecosystems and biodiversity^1,2^. Habitat change and overexploitation are among the human activities with the greatest impacts on vertebrate taxa^3^, and can result in population decline and reduced gene flow, ultimately modifying the evolutionary trajectory of species^3–5^. While animals may initially respond to selective pressures with plastic behavioural adaptations, persistent pressure can drive evolutionary adaptation through genetically based behavioural modifications^6^. This process would drive a population-wide, coherent shift in a series of correlated reaction norms to better align with the environment^7^. Indeed, wildlife can respond to anthropogenic disturbance by adjusting its behaviour to facilitate the coexistence with humans via human avoidance, limited dispersion, habituation − the learning process by which an animal comes to perceive the human presence as neutral − and possibly less aggressive behaviour^8–11^, which has an hereditary component^12^.

The study of the ecology of animals living in human-dominated landscapes has already produced a substantial body of knowledge^6,13–15^, with many reports of behavioural changes^16–18^. For example, Martínez-Abraín & Oro^19^ hypothesised that the prolonged human persecution of several bird and mammal species led to the survival of the shy and less exploratory individuals in a population. However, few studies have linked such altered phenotypes to a genetic base (e.g. ref. ^20^). One great challenge remains to understand the relative roles of phenotypic plasticity and adaptive evolution in response to anthropogenic disturbance in a rapidly changing world^21–23^.

The Apennine brown bear (henceforth called ABB, *Ursus arctus marsicanus*) is a small, endemic and isolated bear population found only in Central Italy, with a long evolutionary history of coexistence with humans. Previous studies using re-sequenced whole genome data showed that this population diverged from other European brown bears 2000-3000 years ago and have remained completely isolated for at least the past ∼1500 years^24^. One of the main factors promoting its isolation was forest clearance and land cover change related to spread of agriculture and increasing human density in Central Italy. Human persecution has been well documented since Roman times^25^, aimed at controlling bear density and expanding urbanised areas^26,27^.

The present ABB population of approximately 50 individuals^28^ shows phenotypic differences from other brown bear populations, such as smaller body size, unique cranial morphology^29–31^, and less aggressive behaviour^24,32^ compared with European, North American, and Asian populations^33,34^. Despite the overall low genomic variation and high levels of inbreeding, Benazzo et al.^24^ showed that high variation was maintained at key loci for survival. This suggests that natural selection might have contributed to shaping the genetic diversity of ABB particularly in cases where adaptation is of critical importance. Interestingly, Benazzo et al.^24^ also found that a pool of genes frequently associated with aggressiveness or tameness in other species was collectively enriched for fixed alternative alleles in ABB, suggesting that behavioural differences with other brown bear populations might be genetically determined. Whether directional selection induced by human persecution targeting more aggressive animals or genetic drift drove this differentiation remains an open question.

Here we focused on the recent evolutionary changes produced by the human activities on this isolated and endangered group of bears. We generated a new high-quality chromosome-level reference genome for the ABB and resequenced 20 whole-genomes of the ABB and bears from a larger European population in Slovakia (SBB), used here for comparison together with published genomes of American brown bears. Population genomic analyses were used to characterize the demographic and selective patterns, with a special focus on the genomic signatures of directional selection in the ABB population. Our findings suggest that selection on genetic variants related to behaviour, likely driven by the human removal of more aggressive individuals, favoured the emergence of a much less aggressive bear population and a less conflictual coexistence between humans and bears.

## RESULTS

### Chromosome-level genome assembly of the Apennine brown bear

Considering the isolation of the ABB from all the other brown bear populations and subspecies, and its unique genetic and phenotypic characteristics, we first assembled an ABB reference genome. We generated 91 Gb of PacBio long-reads and 74.3 Gb of Illumina reads for the chromatin conformation capture (Hi-C) library, corresponding to a sequencing depth of approximately 40X and 34X, respectively. Combining PacBio and Hi-C data, we obtained a primary genome assembly (including the mitochondrial sequence) spanning 2.28 Gb organised in 371 scaffolds, with a N50 of 71.43 Mb and a L50 of 13, in agreement with the genome size estimation from kmers (Supplementary Fig. 1) and the grizzly bear genome assembly^35^. The Hi-C contact matrix supported the presence of 37 scaffolds constituting 99.45% of the assembly (Supplementary Fig. 2), concordant with the known karyotype of the sequenced female (36 autosomes and the X chromosome) (Supplementary Fig. 3). Metrics computed on the final assembled genome suggested a high completeness for the genic and non-genic part of the genome and a low amount of nucleotide errors (Supplementary Data 1).

A total of 22,608 protein-coding genes were predicted and functionally annotated in the ABB genome, showing high completeness metrics, similar to those observed in the grizzly and polar bear genomes (Supplementary Data 2).

When compared with other carnivora species, the ABB genome, included within the brown bear clade in the Ursidae family, confirmed the high synteny levels of all brown bears^36^ while also revealing some chromosomal rearrangements (Fig. 1 and Supplementary Fig. 4). In particular, scanning the ABB genome at a fine scale, we identified a 7.3 Mb chromosomal inversion containing the MHC locus, located at one end of scaffold 31 (Fig. 1). Using a combination of targeted PCR amplification using newly designed primers and long reads mapping, we validated the inversion in four ABB samples, two SBB samples, and one sample from the Italian Alps. This indicates a shared genomic organization among European brown bears, which was not observed in the North American grizzly bear genome. This observation is in agreement with the genomic and mitochondrial differentiation observed between North American and European brown bears^37^.

**Figure 1:**
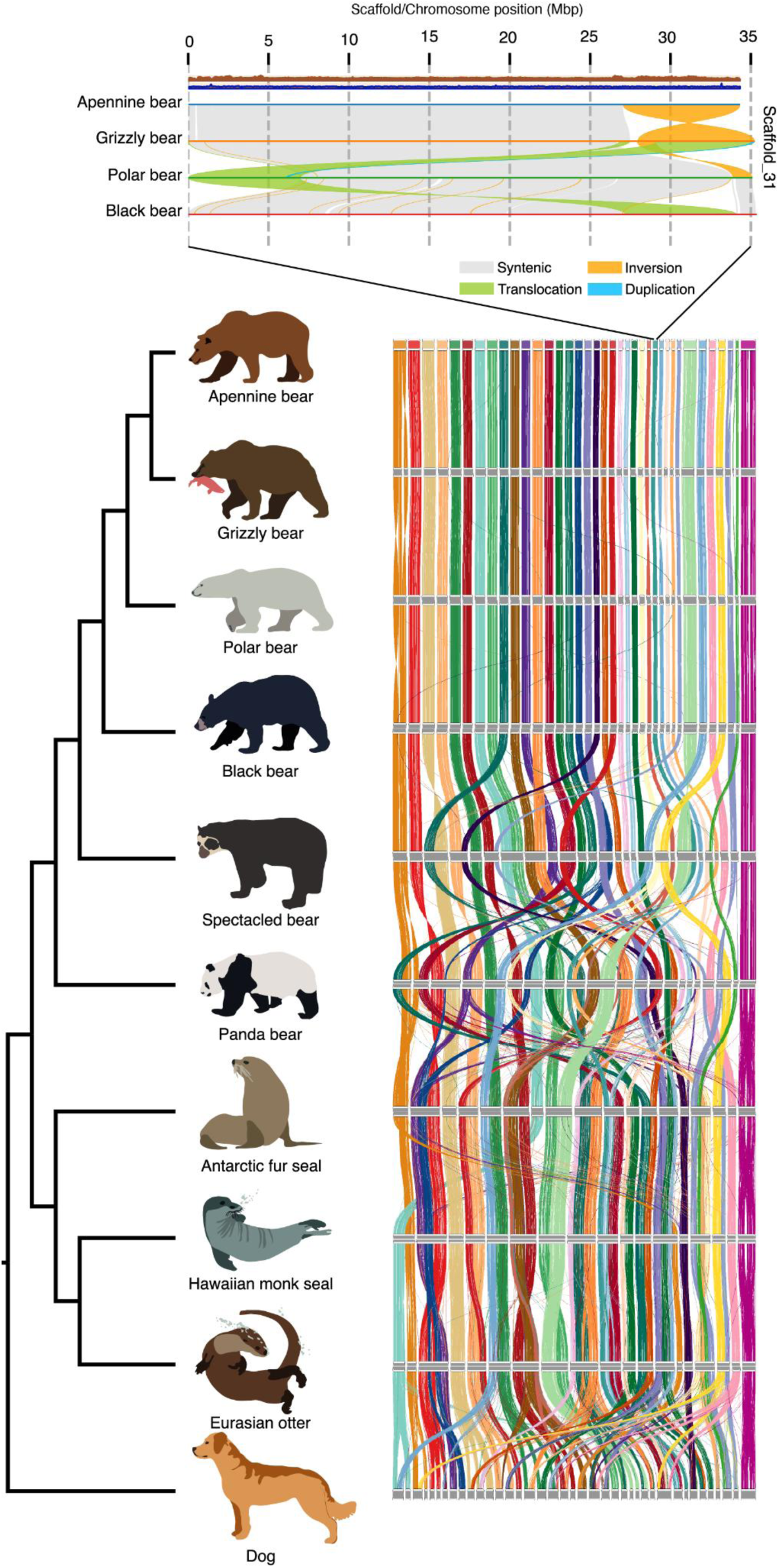
Patterns of chromosome synteny within Carnivora. Pairwise macrosynteny based on 9,054 single copy BUSCO genes. Colours codes are based on ABB scaffold/chromosome organisation. Chromosome alignment between *Ursus* species depicting the MHC inversion between Apennine and Grizzly bears is shown on top. Coverage variations along the scaffold for Illumina and Pacbio reads are also shown in brown and blue tracks, respectively.

Therefore, European bears possess unique chromosomal features, highlighting the need for a distinct reference genome to better study the evolution of this species. Interestingly, the genomic region located at the end of scaffold 31 is also characterised by a chromosome translocation in the polar bear genome, suggesting that this fast-evolving locus might play a key role in the process of species and population divergence^38^.

### Human-driven genomic erosion in the Apennine brown bear

Thirteen high-coverage whole-genome resequencing data from ABB individuals (12 from this study and one from Benazzo et al.^24^) were analysed alongside nine SBB genomes (8 from this study and one from Benazzo et al.^24^) as representatives of the western Eurasian clade of European brown bears^37,39^. Additional genomes from other European bears, available in GenBank (Supplementary Data 3), were used in some analyses, although their sample sizes per population were very limited. Using a multi-reference alignment pipeline, we retrieved 12 million biallelic single nucleotide polymorphisms (SNPs) in the whole dataset, with 2,196,467 and 7,016,199 variants segregating in the ABB and SBB populations, respectively (Supplementary Data 4).

Phylogenetic analysis of the brown bear populations across Europe confirmed that the ABB forms a distinct subclade within the brown bear clade^24^. This subclade is characterised by short terminal branches (Supplementary Fig. 5), in agreement with the lower genomic diversity and higher inbreeding that we found in the ABB compared to other European brown bears (Supplementary Fig. 6A). In particular, all ABB individuals had a Watterson’s theta lower than the one of any SBB individual (Fig. 2a). The SBB had similar genetic variation than bears from all the other European areas, with the exception of brown bears from Spain showing an intermediate value (Supplementary Fig. 6A). Considering regions outside the runs of homozygosity (ROHs), levels of genetic diversity were similar amongst all European bears, suggesting that after excluding the high inbreeding levels in the ABB population (represented by ROH regions), ABB is as diverse as all European brown bears (Supplementary Fig. 6B).

**Figure 2:**
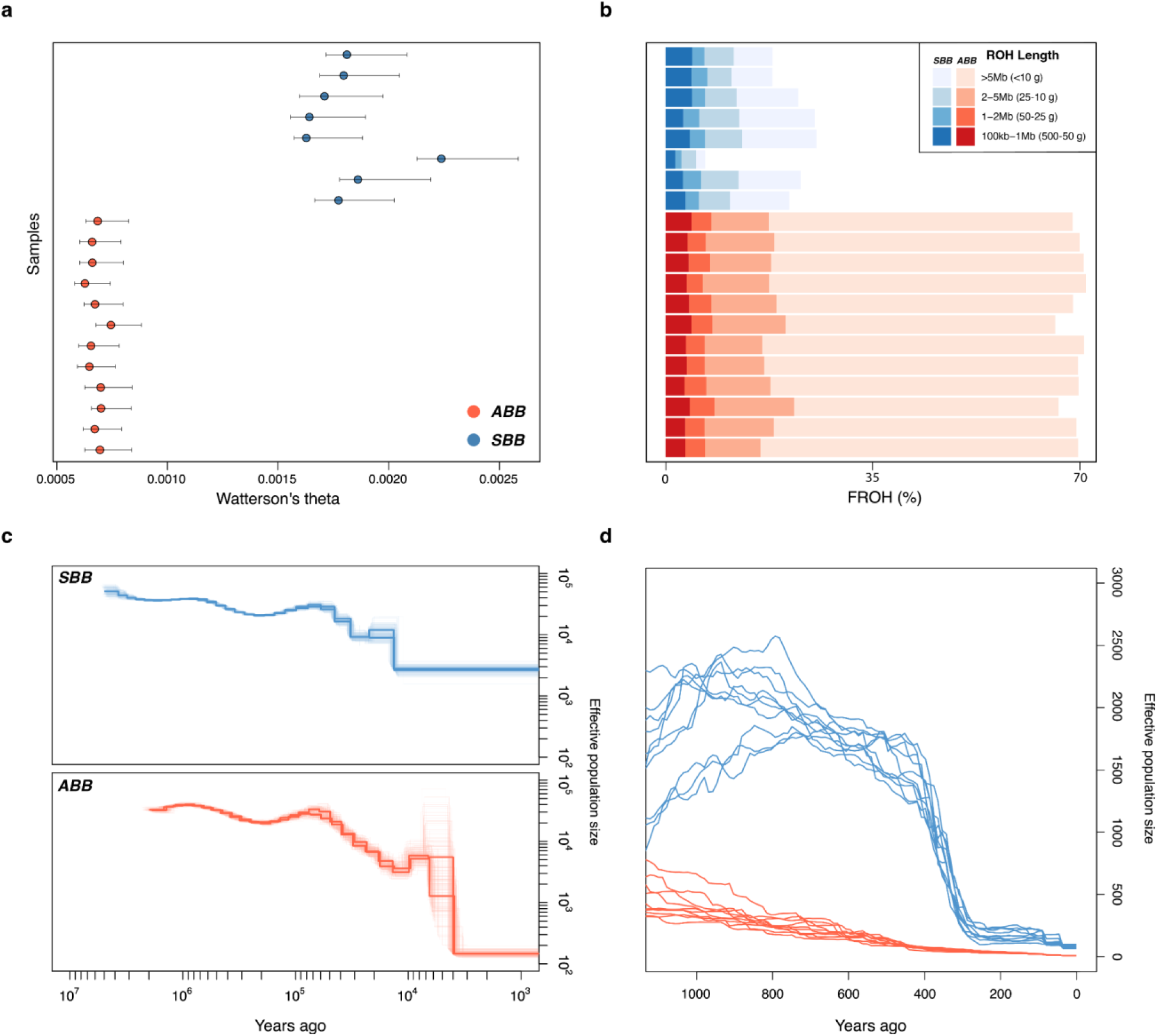
Genomic diversity and demographic history of Apennine brown bears and Slovak brown bears. **a**, Genomic diversity estimates based on Watterson’s Theta. **b**, Inbreeding coefficients (F_ROH_) and corresponding time of inbreeding (generations before present) for each sample. The expected inbreeding time of a particular length of ROH was estimated using g = 100/(2rL), where g is the expected generations that dates back to when both paternal and maternal lineages shared a common ancestor, r is the recombination rate and L is the length of ROH. **c**, Past effective population size dynamics of ABB (8657, 4573 and 4212 samples) and SBB (U1897, U1916 and U1919 samples) based on PSMC. The six genomes with the highest coverage for each population are shown. **d**, Recent effective population size dynamics analysis based on GONE. Ten independent replicates are shown.

The fraction of the genome in runs of homozygosity (F_ROH_) was always higher than 66% in the ABB individuals, whereas in the SBB this fraction ranged from 6.9% to 26% (Fig. 2b). Additionally, all ABB individuals exhibited a higher proportion of very long ROH segments (longer than 5 Mb), likely reflecting recent inbreeding events that occurred within the past 10 generations.

As previously inferred from the analysis of a single individual^24^, the PSMC demographic reconstruction indicated a strong bottleneck in the ABB a few thousand years ago, which was not observed in the SBB (Fig. 2c). This bottleneck, likely driven by deforestation, habitat fragmentation, and land cover changes associated with the introduction and spread of agricultural technologies in certain European regions^40–43^, is the most plausible explanation for the observed differences in genetic diversity between the ABB and other European brown bear populations. Interestingly, when the linkage disequilibrium pattern was used to infer the more recent demographic history over the last millennium, a strong but more recent bottleneck is suggested to occur in the SBB as well (Fig. 2d). Although artefacts due to changes in migration rates through time or to cryptic population structure^44^ could affect the reconstructed demographic history, this recent decline may be linked to the widespread bear eradication policies of the Middle Ages and/or more specifically to the Wallachian colonisation. This refers to the settlement of the Carpathian mountain areas, initiated at the end of the 12^th^ century by the Vlachs, a pastoral ethnic group of Eastern Romance origin, who relied on sheep and goat herding for subsistence^45^.

ABB and SBB showed very similar total numbers of mutations predicted to be mildly and highly deleterious (Fig. 3b and Fig. 3c). However, in SBB, approximately one-third of the sites carrying these mutations were homozygous, while two-thirds were heterozygous. In contrast, the opposite was true for ABB. If we reasonably assume that most deleterious mutations have dominance coefficients close to zero^46^, this result implies that ABB individuals have genomes carrying twice the realized load compared to SBB individuals. However, future consanguineous mating would unmask more deleterious mutations in SBB than in ABB. Considering that a very similar pattern is observed at synonymous sites (Fig. 3a), we should conservatively conclude that the differing patterns of genetic load at SNPs in ABB and SBB are largely driven by their distinct demographic trajectories.

**Figure 3:**
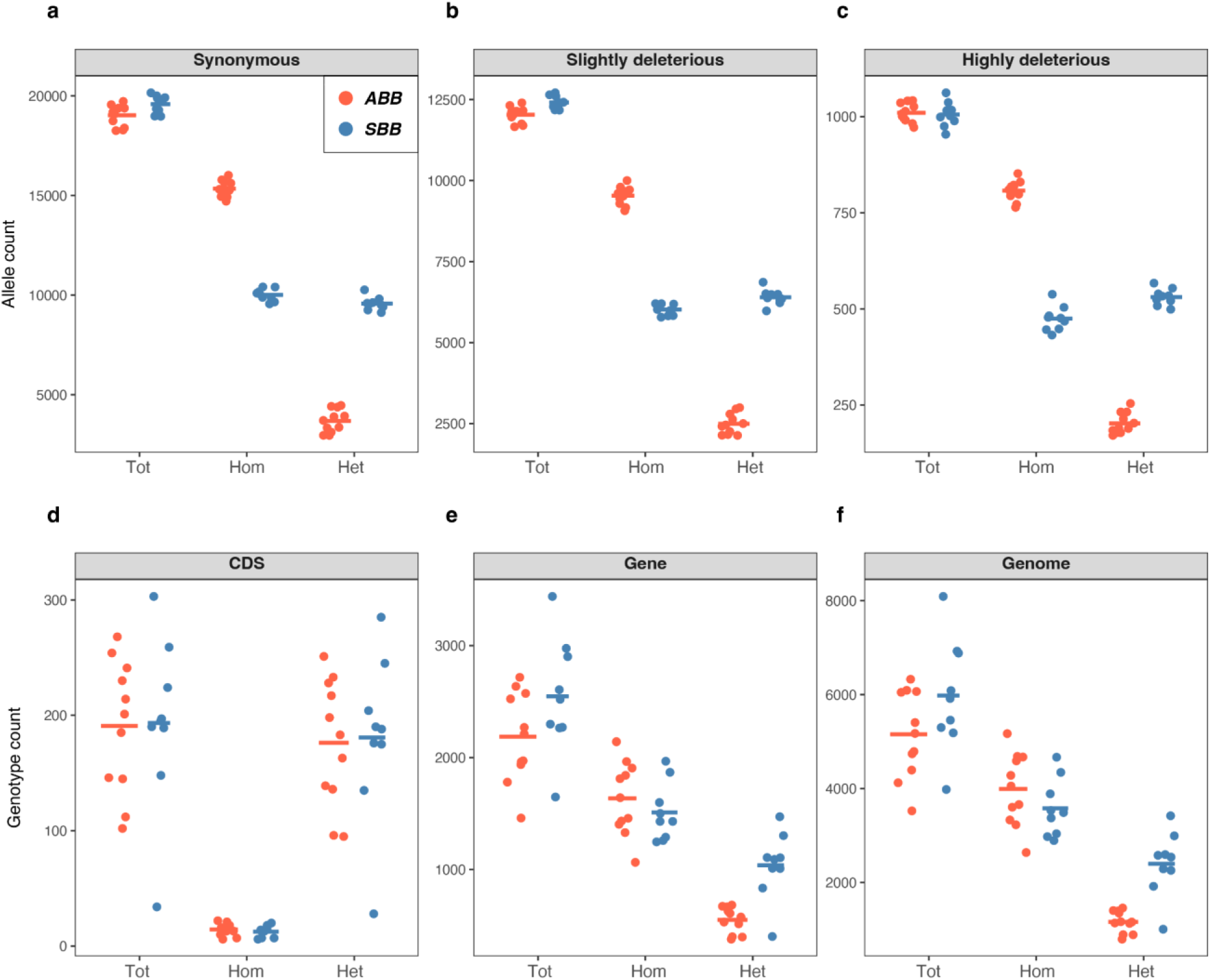
Genetic load comparison of derived variants between ABB and SBB populations. **a**, **b**, and **c** represent SNP-based genetic load categorized as synonymous, slightly deleterious, and highly deleterious, respectively, classified into total (Tot), homozygous (Hom), and heterozygous (Het) variants. **d**, **e**, and **f** show the genetic load measured from the count of large deletions (structural variants between 50 bp and 10 Mb), in coding sequence (CDS), gene-level, and whole genome, respectively, classified into total (Tot), homozygous (Hom), and heterozygous (Het) variants. Horizontal bars indicate the mean of each group.

The estimation of allele age for genome-wide deleterious variants showed that the age of a large fraction of slightly and highly deleterious mutations observed in the ABB population predates the ABB isolation from the European brown bear population (Supplementary Fig. 7). This observation suggests that ABB was connected to the large west Eurasian population identified in de Jong et al.^37^, that acted as a shared reservoir of deleterious mutations that subsequently drifted in ABB contributing to its large realized genetic load.

We also identified high-quality genotyped deletions (DEL) for each individual of the ABB and SBB populations. The DELs were between 50 bp and 10 Mb long. The variation patterns in numbers of DELs across populations (ABB *vs*. SBB), genotypes (homozygote *vs*. heterozygote) and genomic regions (whole genomes *vs*. genes *vs*. coding sequences) are complex and probably affected by several biological and technical factors.

Firstly, the number of DELs observed in coding sequences (CDS) was similar in ABB and SBB individuals, considering the total number or partitioning it in homozygous and heterozygous genotypes (Fig. 3d; Mann-Whitney U test, p-value=0.882, 0.445, and 0.882, respectively). Also, in both populations the majority of DELs in CDS were found in the heterozygous state, with a very low number of homozygous genotypes (Fig. 3d). This pattern, in contrast to what was observed for DELs in other genomic regions (Fig. 3e,f) and across all SNPs categories (Fig. 3a-c), suggests that only a very small number of DELs can be tolerated in homozygosis within the CDS in both populations.

Secondly, we found no significant differences for ABB and SBB in the total amount of DELs per individual at the genomic and genic level (Mann-Whitney U test, p-value=0.152 and 0.131, respectively) (Figure 3e,f; Supplementary Data 5). This pattern was mainly sustained by the derived homozygous DELs (p-value=0.201 and 0.362). Nevertheless, heterozygous DELs were significantly lower in ABB than SBB (p-value=0.0008 at both the genomic and genic level), consistent with their lower genetic variability (Figure 2a). The higher number of homozygous than heterozygous DELs, not expected in the larger SBB population, might be explained by biological or technical factors. In particular, SNPs and DELs might show opposite patterns on genomic diversity, as previously observed in another study^47^. From a technical perspective, the easier identification of homozygous DELs than hemyzygous genotypes^48,49^ could have led to an inflation of the former genotypic state in both populations.

Finally, assuming that gene connectivity, measured by the number of protein-protein interactions, serves as a proxy for a gene’s functional importance^50,51^, we also observed that in all the individuals, genes with DEL alleles in the CDS are, on average, less functionally relevant than genes with DEL alleles in introns (Mann-Whitney U test, p-value=3.26×10⁻⁵, Supplementary Data 6). This finding suggests that deletions in the CDS, which are expected to have a significant impact on the gene product, are more frequently purged in the most essential genes. It also supports our previous result that homozygous DEL genotypes are maintained at tolerable thresholds in both ABB and SBB populations.

### Behavioural shifts in a human-dominated landscape

A previous study revealed that the ABB exhibited a higher-than-expected divergence from other European bears at 22 candidate genes associated with tame/aggressive behaviour^24^. This finding suggested that the ABB could have undergone a genetically mediated shift in behaviour, possibly driven by either genetic drift in its small and isolated population or by the selective hunting of more aggressive or bold individuals during long-term human-bear coexistence in the Apennines. Here, we expanded on this analysis by conducting a blind genome-wide screening for specific signatures of local positive selection and by focusing on genes under putative selection and their functions.

We initially applied different methods specifically designed to detect signatures of positive selection to the SNP dataset: RAiSD^52^, XP-CLR^53^ (Chen et al., 2010), and PBS^54^. Only genes ranking in the top 1% in ABB and not displaying a similar outlier pattern in other populations or population pairs were selected (see Methods for details). A similar approach was applied to the structural variant (SV) datasets; however, only the PBS method was applicable. This first analysis identified a list of 566 genes under selection in the SNP dataset (Fig. 4 and Supplementary Data 7) and 206 genes in the SV dataset (Supplementary Data 8). Twenty genes were common to both lists. To verify that the genes putatively under positive selection exhibited a different variation pattern than the rest of the genome, we computed Tajima’s D statistics for various partitions of the SNP dataset (Supplementary Fig. 8). In both ABB and SBB, random genes and intergenic regions showed an average Tajima’s D between 1.1 and 1.5, respectively, consistent with theoretical expectations for declining populations. For the candidate genes under selection in ABB, the average Tajima’s D was −0.03. Therefore, the putative process of positive selection on these genes shifted the background value toward negative values, as theoretically expected. In contrast, the average Tajima’s D for the same candidate genes in SBB was 1.2, providing further support for the identification of genes uniquely varying in ABB.

**Figure 4:**
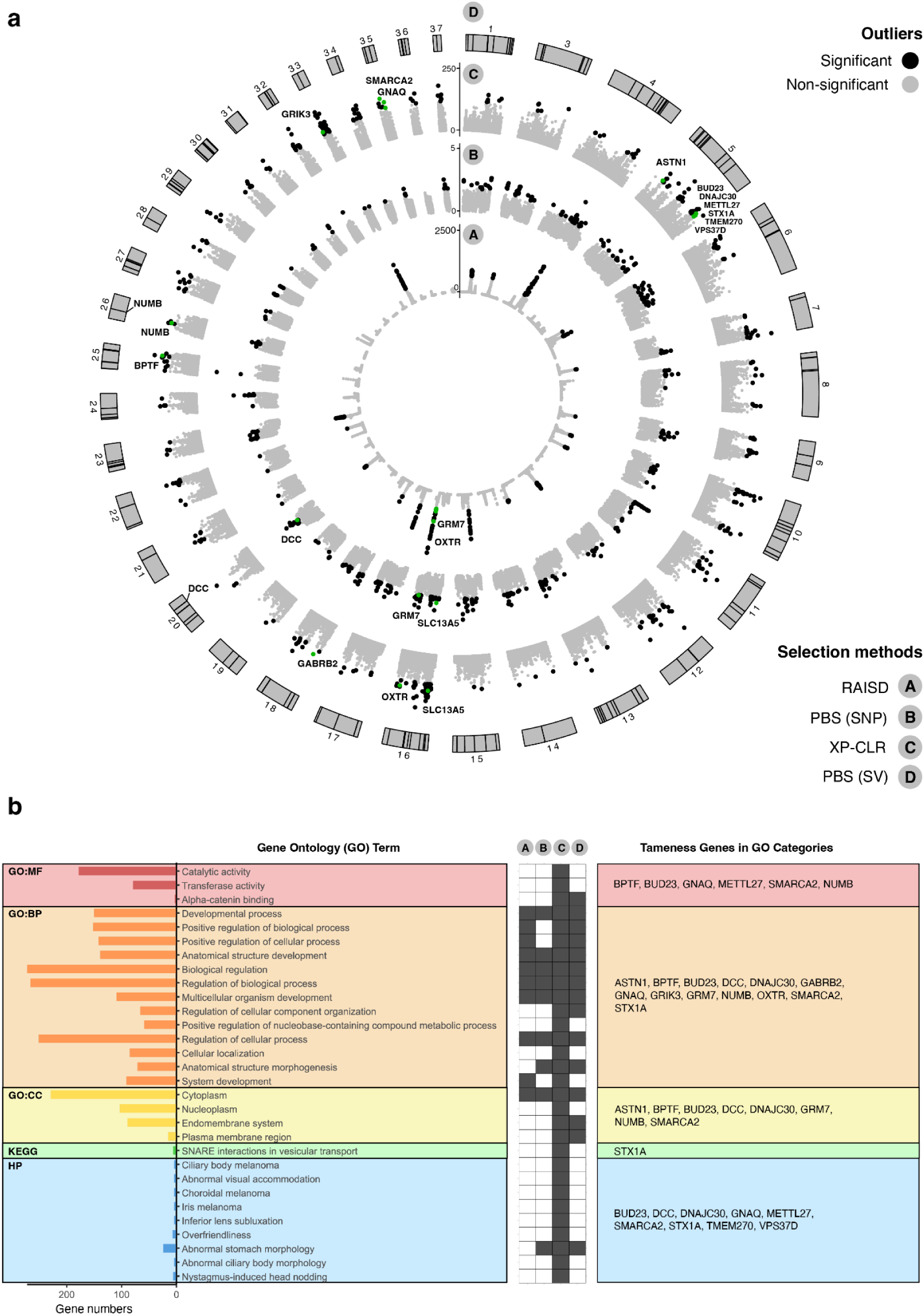
Selection scans and enrichment analyses. **a**, Circos plots of the three selection scans performed on the SNPs dataset (RAiSD, XP-CLR, PBS-SNP) and the selection scan performed on the structural variants (SVs) dataset (PBS-SV) with the Apennine brown bear (ABB) as the target population. RAiSD summarises multiple signatures of selective sweeps in the μ statistic, computed on a single population. XP-CLR compares allelic frequencies between pairs of populations (ABB and SBB in the figure). PBS was performed on four populations, with the ABB as the target. The threshold for defining outlier genomic windows or SVs was the top 1% of the distribution of values for each statistic (black dots). The 17 candidate genes that we identified as possibly determining a less aggressive behaviour in ABB are reported close to their corresponding outlier window (in green). **b**, Enrichment analysis performed separately on the 566 outlier genes from the selection scans on the SNPs dataset and the 206 outlier genes from the selection scan on the SV dataset are summarized with a focus on the terms that were enriched by at least one of the 17 candidate genes related to a less aggressive behavior. For the complete results refer to Supplementary Data S9 and Supplementary Data S10. The total number of genes enriching each term is reported as well. The grey cells indicate that at least one of the aggressive behavior-related genes listed on the right was identified as outlier by the corresponding selection scan.

Next, we performed Gene Ontology (GO) and Human Phenotype (HP) enrichment analyses to identify overrepresented functions and phenotypes potentially linked to tame or aggressive behaviour in the lists of selected genes. Twenty GO terms were enriched in the SNP dataset (Supplementary Data 9), and four in the SV dataset (Supplementary Data 10). The former list included terms like developmental process (p-value=0.0001) and biological regulation (p-value=0.023), suggesting possible developmental and regulatory alterations in the ABB. These could involve the nervous system that plays a central role in behaviour, as five genes with potential influences on tameness/aggressiveness contributed to the enrichment of these terms: BPTF^55^, DCC^56^, GNAQ^57^, GRIK3^58^, and GRM7^59^. The presence of two genes involved in the glutamate pathway, namely GRIK3 and GRM7, is particularly noteworthy since this neurotransmitter and its receptors have a crucial signalling role in the brain. It was suggested that the glutamate pathway was targeted during the tameness experiment conducted on the silver fox in Russia^60^ and during dog^61^ and cat domestication^56^.

The HP enrichment analysis identified nine significant terms, all derived from the SNP data set. Interestingly, the category ‘overfriendliness’ conveys the idea of a docile attitude. This category included seven genes (BUD23, DNAJC30, METTL27, SMARCA2, STX1A, TMEM270, VPS37D), six of which cluster in a chromosomal region deleted in individuals affected by Williams-Beuren syndrome. This genetic disorder in humans is characterized by intellectual disability, short stature, and a heightened eagerness to interact with others^62^. Interestingly, other two genes included in the Williams-Beuren syndrome locus were found to contribute to extreme sociability in dogs^63^.

Finally, a manual screening of the selected gene list identified five additional outliers with roles in brain development. These genes have been previously reported in the literature for their potential involvement in the domestication process of various animals: ASTN1^64^, NUMB^64^, GABRB2^65^, OXTR^66,67^, and SLC13A5^68,69^.

In total, 17 genes putatively under selection for less aggressive behaviour were identified (see Tab. 1). The pattern of divergence between ABB and other brown bears at these genes revealed that the SNPs showing the largest differences in allelic frequencies were located outside the coding regions. Considering the potential phenotypic outcomes arising from the maturation of the same pre-RNA into alternative mature mRNA^70–72^, we focused on genes in which intronic deletions and/or splicing anomalies may have contributed to the observed behavioural shift.

**Table 1:** 17 genes putatively under selection and with possible role in the modification of behaviour in the Apennine brown bear (ABB). The genes are reported together with the criterion for their inclusion in this list. The number of highly differentiated SNPs in intronic (int) and exonic (synonymous: syn; non-synonymous: nsyn) regions is reported. Splicing alterations predicted for the highly differentiated SNPs were divided into two different mechanisms involved in pre-mRNA maturation: the gain or loss of donor or acceptor sites for intron excision and the modification of splicing factor binding motifs. Note that sequence motifs recognized by different splicing factors can be similar and overlapping, hence one SNP can cause the modification or disruption of more than one splicing factor binding site. This analysis was conducted only on the most promising genes in terms of their function and differentiation between ABB and the other brown bears.

| Gene | Criterion | # highly differentiated SNPs int/syn/nsyn | Donor/acceptor splicing site gain/loss | Modifications to splicing factor binding motifs |
| --- | --- | --- | --- | --- |
| ASTN1 | other neural functions | 37/0/0 | 0 | n.a. |
| BPTF | tame list | 3/0/0 | 0 | 3 |
| BUD23 | enrichment | 1/0/0 | 0 | 1 |
| DCC | tame list | 61/0/0 | 0 | 223 |
| DNAJC30 | enrichment | 0/0/0 | 0 | n.a. |
| GABRB2 | other neural functions | 3/0/0 | 0 | n.a. |
| GNAQ | tame list | 86/0/0 | 0 | 65 |
| GRIK3 | tame list | 122/1/0 | 0 | n.a. |
| GRM7 | tame list | 409/0/0 | 0 | 311 |
| METTL27 | enrichment | 1/0/0 | 0 | 1 |
| NUMB | other neural functions | 4/0/0 | 0 | n.a. |
| OXTR | tame list | 34/0/0 | 0 | 26 |
| SLC13A5 | other neural functions | 9/1/0 | 1 | 10 |
| SMARCA2 | enrichment | 72/1/0 | 1 | 62 |
| STX1A | enrichment | 3/0/0 | 0 | 2 |
| TMEM270 | enrichment | 0/0/0 | 0 | n.a. |
| VPS37D | enrichment | 0/0/0 | 0 | n.a. |

In the ABB genome, the genes NUMB and DCC showed signatures of positive selection, supported both by the SNP variant dataset (Supplementary Data S7) and by the presence of an intronic deletion (Supplementary Data S8). NUMB encodes an endocytic adaptor protein and its potential role in the ABB behavioural shift is supported by its involvement in neurogenesis. Notably, an intronic variant in the human ortholog has been associated with bipolar disorder^73^. According to the Mouse Genome Informatics portal^74^ (http://www.informatics.jax.org accessed in April 2024), NUMB is linked to several phenotypes involving abnormal forebrain development. Additionally, it was identified as a target of selection during horse domestication^64^. DCC is a netrine 1 receptor and is involved in neural crest cell migration. It was suggested to directly contribute to the development of tame behaviours in various domesticated animals^56,57^. Interestingly, DCC was also identified as a candidate gene in our study due to a significant allele frequency difference between ABB and other bears at a SNP predicted to be a high-confidence splice-altering variant, leading to the loss of an acceptor site in intron 12 (delta score > 0.5 in SpliceAI analysis).

Another gene of particular interest is SLC13A5, a sodium-dependent citrate cotransporter. This gene has been previously linked to a genomic region associated with temperament at weaning in cattle, a measure of an animal’s docility or unruliness in response to human interaction^68^. Additionally, SLC13A5 has been associated with several human phenotypes related to intellectual disorders, including autism^69^. The potential role of non-coding variants as behavioural modifiers in this gene was supported by the analysis of highly differentiated SNPs. A nearly fixed synonymous mutation in ABB (Scaffold_16:11681411) was linked to an exonic splicing silencer. Specifically, the ratio of exonic splicing silencers (ESS) to exonic splicing enhancers (ESE) was 5 in the sequence context of the major allele in ABB, while it was null for the minor allele according to HOT-SKIP analysis, suggesting that this variant could promote exon 6 skipping during mRNA splicing in ABB. Furthermore, this gene harbours one highly differentiated SNP that was predicted to create a donor splice site according to SpliceAI analysis, and 10 highly differentiated SNPs that were predicted to change the motifs recognized by various splicing factors as suggested by the SpliceAid2 analysis.

SLC13A5, along with DCC, GNAQ, GRM7, SMARCA2, and STX1A, constitutes a set of candidate genes in which ABB exhibit high divergence at SNPs predicted to alter splicing factor binding motifs, including those with known roles in the brain (Supplementary Data 11). Specifically, the splicing factors NOVA1 and NOVA2^75^ have been implicated in the evolution of alternative splicing patterns in the brain across vertebrates, primarily through modifications of their splicing enhancers and silencers^76^. Of particular interest, the candidate gene DCC stands out as a known NOVA2 target^77^.

## DISCUSSION

Understanding how wild species respond to anthropogenic pressure over time is crucial for their preservation and successful coexistence with humans. In this study, both previously published and new whole genome sequencing data (including a new chromosome-scale reference genome) were used to investigate the genomic impacts of long-lasting persecution and isolation in a small relict population of brown bears in Central Italy, where cohabitation with local human communities have been documented for several millennia. Low genomic variation, high levels of inbreeding and a substantial amount of realized genetic load characterize this population, likely as a consequence of random drift effects during a human-induced demographic reduction.

Different methods, however, also point to genes affected by positive selection processes specifically occurring in the ABB. Our results show that some variants associated with these genes, likely present as standing variation in regulatory regions before the selection occurred, might be responsible for the docile phenotype observed in these bears. The most plausible explanation for the spread of these traits is selective culling. In other words, despite the capacity of the brown bear to implement plastic behaviours to cope with anthropogenic disturbance^17^, our data support the hypothesis proposed by Festa-Bianchet^78^ and by Allendorf & Hard^79^ for harvested wild populations – that is, the more aggressive bears were eliminated, favouring the spread of genetic variants associated to reduced aggressiveness.

Wild populations that are able to persist in human-dominated landscapes, especially those heavily impacted by overexploitation, deserve special attention in order to study, understand, and preserve adaptations that might mitigate conflicts and promote coexistence^8,10^. In particular, the reduced aggressiveness of the ABB, possibly due to the spread of alleles that control genetically this behavioural trait, has lowered the negative attitude towards bears in the human population that cohabits with this large mammal^80,81^. This, in turn, can reduce the human-caused mortality of the species. The ultimate result is the increase in the chances of population persistence in a human-dominated environment, despite an extreme drop of genomic variation and the accumulation of deleterious mutations. Therefore, even when pervasive genomic erosion has impacted a small, isolated wild population, genetic adaptations can play a crucial role in reducing the extinction risk.

## MATERIAL AND METHODS

### Long-read sequencing

For the genome assembly, PacBio long reads were produced for a single Apennine brown bear (ABB) individual. A blood sample (500 μL) was drawn from a female Apennine bear (Lauretta) kept in captivity in the faunistic park “Centro Visita di Pescasseroli” (L’Aquila, Italy) during routine veterinary checks. The blood sample was immediately frozen in liquid nitrogen to preserve the integrity of nucleic acids. High molecular weight DNA was isolated from 200 μL of blood using the Nanobind Tissue big DNA kit (Circulomics Inc., Baltimore, USA). DNA quality and fragment length were checked in a pulse field gel electrophoresis and DNA concentration was measured with fluorometric and spectrophotometric assays using a Qubit 2.0 Fluorometer (Invitrogen, Carlsbad, California, US) and a TECAN Nanoquant Infinite 200 Pro (Tecan Mannedorf, Switzerland), respectively. Fragments of 35,000 bp length were selected using a Blue Pippin device (Sage Science; Beverly, MA, USA). Isolated fragments were used to prepare the DNA library with a SMRTbell express template prep kit 2.0 (Pacific Biosciences, Menlo Park, California, US) according to manufacturer’s protocols. The library was run on eight PacBio SMRT Cells 1M in continuous long-read sequencing (CLR) mode on a PacBio Sequel platform.

### Resequencing of ABB and SBB samples

For population genomic analyses, a total of 12 and 8 *U. arctos* samples were collected in Central Italy and Slovakia, respectively. Within the 12 ABB, two cubs and their mother were included. Details of the samples are reported in Supplementary Data 3. The samples were then preserved in 96% ethanol at −20 °C and the total-cell DNA was extracted using a PureLink Genomic DNA Mini Kit (Invitrogen). DNA integrity was assessed by 1.5% agarose gel electrophoresis and DNA concentration was measured using a Qubit 4 fluorometer Broad Range Assay (Invitrogen). Short-read genomic libraries were constructed using an Illumina DNA PCR-Free Prep Kit (Illumina) according to the manufacturer’s protocol. Libraries were sequenced paired-end on an Illumina NovaSeq 6000 System using a 300-cycle S2 Reagent Kit v1.5, with a target coverage of 10-15X for all samples.

### Cell cultures, karyotyping and Omni-C

To confirm the chromosomal structure of our assembly, a karyotype for the ABB was generated using a cultured cell protocol. A gingival tissue biopsy was obtained from the bear Lauretta. Cells were cultured in a medium composed of 50% RPMI1640 and 50% Iscove’s Modified Dulbecco’s Medium, supplemented with 10% fetal bovine serum, 1% penicillin (10,000 units/mL) - streptomycin (10 mg/mL), 1% gentamycin sulfate (10 mg/mL), 0.5% amphotericin B (250 mg/mL) and 1% L-glutamine (200 mM) and incubated at 37 °C with 5% CO_2_. Chromosome preparations were made following standard procedure^82^. In brief, after 4 h of treatment in 0.01 ng/mL colcemid, the cells were removed by standard trypsinization and placed in a 15 mL tube. Cells were then centrifuged at 10,000 g, surnatant was removed and substituted with a 1:1 mixture of 0.075M KCl and 0.4% sodium citrate (hypotonic treatment). After a 20 min exposure at 37 °C the cells were pelleted by centrifugation and fixed in methanol:acetic acid fixative (at a ratio of 3:1). Slides were then prepared by dropping metaphases with a Pasteur pipette onto a clean glass microscope slide. Diploid number and chromosome morphology were determined from the analyses of 20 mitotic cells stained with DAPI. Karyotype was arranged according to the standard ursid karyotype set^83^.

Part of the cultured cells were harvested and sent to Dovetail Genomics (Scott’s Valley, CA) to construct chromatin conformation capture libraries using the Omni-C kit from Dovetail Genomics. Omni-C libraries were sequenced paired-end on an Illumina NovaSeq 6000 System using a 300-cycle Reagent Kit v1.5.

### Genome assembly

Raw PacBio CLR reads were screened with FastQC v0.11.9^84^ to detect the presence of residual adapter sequences and possible contaminants and trim the reads accordingly. Reads were then assembled into contigs using CSA v2.6^85^ with a combination of specific flags optimized for reads generated by a Pacbio’s Sequel instrument (-p 0 -k 15 -L5000 -S 2 -A). Before the scaffolding step within the CSA pipeline, the assembled contigs were polished with pbgcpp v2.0 (Pacific Biosciences) and POLCA v4.0^86^. using PacBio’s CLR and Dovetail’s Omni-C data, respectively. One round of purgedups v1.2.5^87^ was performed to remove the residual haplotype redundancy. Omni-C reads were then mapped to the scaffolds and processed according to Dovetail’s documentation (https://omni-c.readthedocs.io/en/latest/index.html) using default parameters and contigs were scaffolded using SALSA v2.3^88^ setting the “-e DNASE” option and skipping the contigs misassembly step. The CSA pipeline was completed using the reference genome of the grizzly bear (GenBank accession: GCA_023065955.2) to further scaffold the genome and improve contiguity. To verify that the scaffolding step using the grizzly bear genome did not introduce any misassembly due to the divergence of the two bears groups, the Omni-C reads were remapped to the final genome and PretextSuite v0.1.8^89^ was used to visually inspect the Hi-C contact maps and to manually anchor/correct scaffolds to maximize the fit of the contact data. Remaining gaps between scaffolds were closed with TGS-GapCloser 1.2.1^90^ using PacBio CLR reads. The mitochondrial sequence was reconstructed using mitoVGP pipeline v2.2^91^ using the grizzly bear mtDNA (GCF_003584765.1) as reference and then it was manually added to the final assembly.

### Transposable Element and Repeat Annotation

To identify and annotate transposable elements in the ABB genome, we initially constructed a *de novo* repeat library using the Extensive de-novo TE Annotator (EDTA) v1.9.9^92^. Subsequently, we refined the library using DeepTE^93^, which employs convolutional neural networks to classify unknown elements at the order and superfamily levels. Then we used RepeatMasker v4.1.2^94^ with the final library to mask the genome and we parsed RepeatMasker output files with RM_TRIPS script (https://github.com/clbutler/RM_TRIPS).

### Gene annotation

RNAseq data from closely related species available on NCBI (see Supplementary Data 12) were used as evidence for the gene prediction. We mapped the quality- and adapter-trimmed RNA-seq reads to the soft-masked assembly with hisat2 v2.1.0^95^ with standard parameters followed by sorting and indexing with samtools v1.10^96^. Quality control and trimming for adapters and low-quality bases (quality score <20) of the RNA-seq raw reads were performed using FastQC and TrimGalore v0.5.0 (https://github.com/FelixKrueger/TrimGalore), respectively. All the BAM files were filtered to remove invalid splice junctions with Portcullis v1.1.2^97^. For the prediction of gene loci and structures, we used the BRAKER3 pipeline v3.0.2^98^, which uses the soft-masked genome, filtered RNA-seq alignments and OrthoDB11 protein data^99^ (https://bioinf.uni-greifswald.de/bioinf/partitioned_odb11/) as input to subsequently run the gene prediction tool GeneMark-ETP v1.00^100^ and then AUGUSTUS v3.4.0^101^ for its annotation process. The BRAKER transcript selector, TSEBRA v1.0.3^102^, was employed to unify predictions from these 3 sources and configured such that RNA evidence had greater weight than protein evidence. The resulting gtf file was converted to a fasta file using the perl script, gtf2aa.pl, which is included with the Augustus programming suite. To assess the completeness of the annotation, we used the Mammals odb10 reference database in protein mode in BUSCO v5.2.2^103^. Sequences were searched in the nonredundant NCBI protein database using Diamond v0.9.10^104^ with an E-value cut-off of ≤1 × 10^−5^. Blast2go v5.0^105^ and InterProScan v2.5.0^106^ were used to assign Gene Ontology (GO) terms. Protein domains were annotated by searching the InterPro v32.0^107^ and Pfam v27.0^108^ databases, using InterProScan v5.52 and HMMER v3.3^109^, respectively.

### Phylogeny

Orthologous groups in Ursidae family genomes were identified from the predicted protein sequences of the ABB and other six bear genomes already published: *U. americanus*^110^, *U. maritimus*^111^, *U. thibetanus thibetanus*^112^, *U. a. horribilis*^113^, *Tremarctos ornatus* (https://www.dnazoo.org/assemblies/tremarctos_ornatusC) and *Ailuropoda melanoleuca*^114^. As outgroups, we included genomes from three Carnivora families (Phocidae, Mustelidae and Canidae): *Neomonachus schauinslandi*^115^, *Lutra lutra*^116^ and *Canis lupus familiaris* (ROS_Cfam_1.0, GCF_014441545.1). We used the longest transcript to represent the gene model when several alternatively spliced transcripts of a gene were annotated. We used a series of tools to cluster proteins into orthogroups, reconstruct gene trees and estimate the species tree: OrthoFinder v2.5.4117, Diamond v0.9.14, Multiple Alignment using Fast Fourier Transform (MAFFT) v7.305118 and RAxML v8.2.12119.

### Synteny

The syntenic relationship within Carnivora was evaluated comparing the ABB genome with the grizzly bear and other eight species having a chromosome-scale reference genome from NCBI or DNAzoo^120,121^ repositories including *U. americanus*, *U. maritimus*, *U. a. horribilis*, *T. ornatus*, *A. melanoleuca*, *N. schauinslandi*, *Arctocephalus gazella*, *L. lutra* and *C. l. familiaris*. BUSCO v5.0.0 was used to scan each genome using the carnivora_odb10 database and all complete single-copy genes shared among species were extracted for comparative analysis. The coordinates of each BUSCO gene was traced between species using the software RIdeogram v0.2.2^122^. Large structural variants characterising the genome of the five bear species were identified and visualised using syri v1.7^123^ and plotsr v1.1^124^, respectively.

### MHC inversion

The presence of a rearrangement on Scaffold_31 was assessed using region-specific amplification followed by Sanger sequencing. Specifically, a primer pair was designed to amplify the region across the putative breakpoint of the scaffold followed by the inverted region based on the latest assembly of the ABB genome. The forward primer annealed 1,066 bp upstream of the breakpoint (Umar_inv_F1 5’-GTCATTCATGGCCAGGGAAC-3’), while the reverse primer annealed 1,922 bp downstream of the breakpoint (Umar_inv_R1 5’-GCTCTGACTCCTTCCACCAT-3’), allowing the amplification of a 2,989 bp fragment. The primer pair was used in a PCR reaction with 1.25 units of the enzyme PrimeSTAR GXL DNA Polymerase (Takara), 30 cycles with annealing at 58°C for 15 seconds and extension at 68°C for 3 minutes. The resulting PCR products were then purified and Sanger sequenced with a combination of internal primers (Umar_inv_R2 5’-CATAACCACACAGCCAGCAA-3’; Umar_inv_R3 5’-CATGCTGTCATCCCCAAAGG-3’; Umar_inv_F3 5’-GCTTAACCAACTGAGCCACC-3’) to confirm the inversion. The analysis was carried out on four ABB including Lauretta, two samples from Slovakia and one sample from the Italian Alps. To further validate the inversion, the read mapping distribution along the region of Scaffold_31 was generated aligning short- and long-reads to the reference genome using bwa v0.7.17 and minimap2 v2.22 respectively. The genomic tracks showing the sequencing coverage were generated from the alignments using plotsr v1.1.

### SNP variant calling

For population-level brown bears samples, we used the newly produced reference genome of the ABB for mapping: 20 individuals resequenced for the purpose of this study (including 12 ABB and 8 SBB), and 9 brown bear individuals from Europe already available (including 1 ABB and 1 SBB)^24,125,126^. We also mapped three polar bear individuals^125^ to the polar bear genome (GCF_017311325.1: 3900 scaffolds, total length=2,330,485,043 bp) and three black bear individuals^127–129^ to the black bear genome (GCF_020975775.1: 105 scaffolds longer than 50,000 bp and covering 2,328,269,719 bp, i.e., 90% of the genome length was used). All reads were mapped to the corresponding reference genome using bwa-mem v0.7.17^96^ with default parameters, with an Optical Density (OD) value of 2,500 for the samples generated in our study that used patterned flow cells and 100 for the other individuals. Reads were merged prior to mapping using AdapterRemoval v2.3.1^130^ when the sequencing strategy generated overlapping reads. PCR and optical duplicates were tagged using MarkDuplicate from Picard v2.24.1 (http://broadinstitute.github.io/picard). For the individuals sequenced in this study, a PCR-free sequencing kit was used so reads tagged as PCR duplicates were further untagged using a custom bash script. Diverse statistics were computed to ensure the quality of the mapping.

An independent SNP calling was performed for each of the three species. SNP calling was performed using GATK v4.1.9.0^131^ following the best practice guides for short variant discovery, composed of three steps: calling variants per sample using HaplotypeCaller; consolidate GVCFs to improve scalability using GenomicsDBImport; joint-genotype all the per-sample GVCFs using GenotypeGVCFs. HaplotypeCaller was run independently in each scaffold of each individual, using the option EMIT_ALL_CONFIDENT_SITES to output all sites (including non-variant sites) and the output were generated in condensed non-variant blocks. A minimum mapping quality of 20 was used. GenomicsDBImport and GenotypeGVCFs were run independently for each scaffold. The tag --include-non-variant-sites was used in GenotypeGVCFs to output all sites (including non-variant sites). For the brown bears, all scaffolds were cut into windows of 10 Mb using bedtools v2.30 makewindow^132^, and each window was run independently to satisfy memory requirements. Finally, all the windows and scaffolds were merged to produce one unique VCF file per dataset using bcftools merge v1.11^133^.

From the raw VCF file produced after SNP calling, we first extracted SNPs and indels using GATK SelectVariant. We then filtered indels by quality (> 60) and then filtered out SNPs that were located within 5 bp of the high quality indels, and that fulfilled the following criteria: QUAL < 60; QD < 2.0; FS > 60.0; MQ < 40.0; MQRankSum < −20.0; ReadPosRankSum < −8.0; DP < mean depth/3; DP > mean depth*2; GQ < 10. The mean depth across individuals was computed with samtools v1.11 at each position. We also extracted invariant sites using GATK SelectVariant and further filtered them using GATK VariantFiltration to remove genotypes with RGQ<10. We then merged high quality variant and invariant sites using bcftools concat.

The three reference genomes used for read mapping were aligned to identify syntenic regions that could be safely associated and used in every analysis involving a comparison between species. The aim of this procedure was to remove the reference bias associated with short-read mapping toward a single reference and to exclude variants located within, or in proximity of, genomic regions involved in cross-species rearrangements. The polar and the black bear genomes were aligned to the ABB reference using minimap2 v2.22^134^ setting a maximum divergence of 5% and adding the “cs” tag. Transanno v0.2.4 (https://github.com/informationsea/transanno) was then used to produce a chain file from minimap2 output, to extract well aligned regions between reference genomes and to lift the genomic coordinates of all sites (monomorphic and polymorphic) contained in the VCF files between species. The lifted VCF files for each species, all of them following the ABB genome coordinate system, were merged using bcftools, retaining only sites for which the liftover procedure between species was possible and excluding sites completely missing in all individuals belonging to one or more species.

### Structural variants calling

We performed the Structural Variant (SV) calling on the genomes of 11 ABB from Central Italy (mother-daughter pair excluded), 9 SBB, 8 grizzly bears from Canada and 6 grizzly bears from Alaska. To avoid reference bias, all reads were mapped to the polar bear (*U. maritimus*) reference genome using the same procedure detailed for SNP calling. To perform the SV calling we used the meta-method Parliament2 v0.1.11^135^ that incorporates multiple pre-installed SV callers to identify high-quality SVs starting from short-reads genomic data. Of the different pre-installed SV callers we selected BreakDancer^136^, CNVnator^137^, Lumpy^138^, Delly^139^ and Manta^140^ in order to detect different types of SVs (deletions, duplications, insertions, and inversions) using different signals (paired-end reads, split reads and read depth). To identify the SVs in each individual’s genome, Parliament2 requires the BAM file and the reference genome used for the alignment. The SV calling was limited to scaffolds longer than 1 Mb with the option --filter_short_contigs. Next we used SURVIVOR v1.0.6^141^, in its option “merge”, to obtain a consensus of the SVs identified by each method. Parameters have been set to maintain only SVs longer than 50 bp called by at least 2 SV callers out of 5 and to group as single SV those with the same type and a maximum difference of 1000 bp between starting and ending positions without considering the genotype. Then, SURVIVOR merge was used to obtain a multi-sample VCF with all the SVs identified in at least 1 individual out of 35. Then we used vcftools v0.1.16^142^ to filter out SVs with low quality value (GQ < 20) and that were no longer identified in at least 1 individual (flag --max-missing-count). We removed the SVs with a distance between the starting and ending positions longer than 10 Mb and those shared between all individuals. Only for duplications and inversions we chose to also remove the ones partially overlapping in all individuals using bedtools merge and intersect.

### Species phylogeny and genomic diversity

To reconstruct the maternal lineage through mtDNA, we mapped raw reads using the read alignment procedures previously described for the resequencing data. To generate consensus sequences for each male, we used bcftools mpileup v1.10.2-9, filtering out bases with low phred-scaled quality scores (Q < 20) and reads with low mapping quality (Q < 40). An alignment totalling 13,882 bp which included only coding regions and rRNA (12S and 16S) was used in the analysis. The phylogeny was reconstructed using a neighbor-joining tree implemented in Geneious 8.1^143^ with GTR+alpha substitution model. The black bear sequence was used to root the tree.

To analyse the DNA nuclear phylogeny, we computed pairwise genetic distances between individuals using the function --distance square 1-ibs implemented in PLINK v1.90^144^ for 34 individuals at 24,868,223 positions. We then estimated a neighbour-joining phylogenetic tree from the pairwise distance matrix using the package ape^145^ in R v4.0.

To estimate genomic diversity, we first computed different statistics (number of SNPs and Watterson’s θ) in the studied populations. Then we inferred runs of homozygosity (ROH) with ROHan v1.0^146^. To obtain an estimate of the maximum heterozygosity level permitted in ROH regions, we ran ROHan on the chromosome X for all male individuals, removing the first 11 Mb that exhibited an abnormal high heterozygosity level and excluding repeated regions. We obtained a mean heterozygosity of 1.36e^-4^, and subsequently used a rohmu value (the inferred heterozygosity rate) of 1e^-4^ to define regions in ROH in the autosomes for all ABB and SBB populations. We conducted our analysis using a window size of 100 kb and calculated the fraction of the total genome assigned to runs of homozygosity (F_ROH_), a commonly used proxy for the inbreeding coefficient. Additionally, we estimated the distribution of different ROH lengths from ROHan output files. To estimate the number of generations since inbreeding events, we applied the formula *g* = 100/(2*rL*), where *r* is the recombination rate (1 cM/Mb) and *L* is the ROH length in Mb^147^. Based on this, we estimated that an ROH length of 100 kb corresponds to ∼500 generations, 1 Mb to ∼50 generations, 2 Mb to ∼25 generations, and 5 Mb to ∼10 generations. ROHan was also used to estimate Watterson’s θ both including and excluding ROH regions.

### Demography

The long-term variation in population size was investigated using PSMC v0.6.5^148^ following the standard procedure for the three individuals with highest depth of coverage for ABB and SBB, respectively. SNPs were called from the BAM files for each of the 36 autosomal scaffolds longer than 10 Mb using bcftools mpileup and bcftools call, only using positions with a minimum base and mapping quality of 30 and covered between one third and two times the mean individual depth. Windows of 100 bp were then used to produce the PSMC input. The default time patterning (“4+25*2+4+6”) was used, with an initial theta/rho ratio (-r parameter) of 5 and a maximum 2N_0_ coalescent time (-N parameter) of 15 (default values). We performed 100 rounds of bootstrapping by randomly sampling with replacement from 5 Mb sequences. A generation time of 11 years and a mutation rate of 1.82e^-8^ per site per year^24,125^ were used for scaling.

Since the PSMC does not infer recent *N*_e_ variation (<400 generations), we used a different method to infer recent changes in *N*_e_ that might have influenced the ABB population in comparison to its SBB counterpart. We used GONE v1.0^149^, a method based on linkage disequilibrium. GONE was run for the 11 ABB (excluding the mother-daughter pair) and 9 SBB to estimate recent demographic history in European brown bears. Default parameters were used except for *hc*=0.01^150,151^. We performed 10 independent replicates for each population to evaluate the consistency of the estimations.

### Mutation load

We first polarised each SNP as ancestral or derived using two outgroups: the polar bear (3 individuals) and the black bear (3 individuals). We defined the ancestral allele as the allele present in at least 2 out of 3 polar bear individuals and in at least 2 out of 3 black bear individuals using a custom python script. All sites where at least one of these outgroups was heterozygous were discarded to maximise the confidence in the ancestral allele definition.

We assessed the mutational load in both 11 ABB (mother-daughter pair excluded) and 9 SBB through two distinct methods. We restricted these analyses to the protein coding sequences (CDS), extracting the CDS of the longest transcript of each gene of the annotation and merging the overlapping windows with bedtools, resulting in a total CDS length of 33 Mb. First, we measured the relative mutational load in each individual as the number of derived alleles at sites that are under strict evolutionary constraints (i.e., highly conserved) and thus likely to be deleterious using Genomic Evolutionary Rate Profiling scores (GERP)^151^. GERP scores for the Polar bear in bigwig format was obtained from a multiple alignment with 91 mammal species downloaded from ENSEMBL (https://ftp.ensembl.org/pub/current_compara/conservation_scores/91_mammals.gerp_conservati on_score/). We firstly aligned the ABB assembly to the polar bear assembly using minimap2 with parameter -cx asm20. The GERP scores were subsequently transferred to ABB reference coordinates using Transanno between the polar bear and the ABB genome. Our analysis considered both heterozygous positions (counted as one allele) and homozygous positions (counted as two alleles), recognizing that the mutational impact of heterozygous positions involves additional assumptions regarding the dominance coefficient. GERP identifies constrained elements within multiple alignments by quantifying substitution deficits, reflecting substitutions that would have occurred if the element was neutral DNA but did not due to functional constraints, accounting for phylogenetic divergence. The individual relative mutational load was calculated by summing the number of derived alleles above a GERP score of four (highly deleterious).

Secondly, we run SnpEff v5.0^152^ on the polarised SNPs present in CDS to classify each mutation as synonymous, missense (*i.e.*, non-synonymous) or nonsense (including stop-gained, start-gained, start lost, stop codons, splice donor variant and splice acceptors). For each class of putatively deleterious mutation (missense and nonsense mutations), the genetic load was separated into two components: i) the masked load estimated as the individual number of heterozygous sites, and ii) the realized load estimated as the individual number of homozygous derived sites^153^.

We estimated the age of deleterious and neutral alleles by inferring the time to the most recent common ancestor (TMRCA) of those mutations. We used Genealogical Estimation of Variant Age (GEVA)^154^, which reconstructs a genealogical tree at the chromosomal level of variable sites and estimates the age of genetic variants by identifying segments that are shared between a pair of haplotypes. It assumes that an allele originates from a mutation event on the genome of a common ancestor, and it is subsequently passed down to all descendant haplotypes. The time of origin of a mutation is obtained by a probabilistic estimate that combines the cumulative distributions for pairs of haplotypes that share the mutation (“concordant pairs”) and the pairs that do not (“discordant pairs”). Since GEVA requires phased data, we first phased the SNPs with BEAGLE v5.4^155^, using the options impute=false, Ne=10.000, burnin=15, iteration=40 and phase-states=300. Singletons, private doubletons, and sites with more than 10% of missingness were excluded from the analysis with vcftools. We used an effective population size of 10,000, a recombination rate of 1.0 × 10^-8^ and a mutation rate of 1.82 × 10^-8^ per site per year.

We further investigated the genetic load attributable to the SVs by comparing the amount and distribution of DELs in the ABB and the SBB populations. To obtain the SV genotypes for the ABB and the SBB individuals we first kept only those 20 samples (11 ABB and 9 SBB) from the merged dataset, and selected only DELs as a proxy of the SV dataset being the most abundant SV category (82% on average per individual). To exclude the SVs accumulated from the separation between the brown bears and the polar bear (used as reference genome), we filtered out the SVs with a frequency greater than or equal to 0.85, considering them fixed but not identified in all 20 samples due to methodological limitations in SV identification with low-medium coverage. Starting from the individual genotyped VCFs produced by Lumpy and genotyped by SVTyper^156^ during the SV calling step, we selected only the DELs that were included in the 20 samples merged SV dataset as we considered them as high-quality SVs using a custom script. For each individual we quantified the number of DELs, and the derived homozygous and heterozygous genotypes considering the whole genome, the genes and the CDS. To further analyse the characteristics of genes affected by homozygous and heterozygous DELs, we focused on the number of protein-protein interactions as a proxy for gene importance^50,51^. We downloaded the protein-protein interaction database from STRING^157^, which lists all interactions between proteins in the polar bear, in order to calculate the number of interactions for each gene product with a minimum confidence score of 0.4. We quantified the number of interactions of the gene products of genes impacted by both homozygous and heterozygous DELs in their CDS or in their intronic regions and the mean number of interactions for all the gene products listed by STRING. Then, we used a Mann-Whitney U test to compare these mean numbers of interactions.

### Selection scans

We implemented a series of analyses to detect selection in the ABB. As we expect this small population to have experienced considerable levels of genetic drift, that can leave confounding signatures to the ones of positive selection, we increased the number of populations studied and the number of approaches in the attempt to infer positive selection. We included 15 individuals from grizzly brown bears (9 from Canada^37^, 6 from Alaska^37,126,127^) to compare North American to European populations. For these new samples, we followed the same procedure as described above for read mapping (to the ABB reference genome), SNP calling and filtering including all brown bear individuals. This new dataset comprised 35 individuals (see Supplementary Data 3) and 12,906,067 SNPs.

To detect signatures of recent positive selection at the population level, we used RAiSD v2.9^52^. RAiSD detects positive selection based on multiple signatures of selective sweeps, namely the change in the site frequency spectrum, the levels of linkage disequilibrium and the amount of genetic diversity along a chromosome, scoring these three components into the *μ statistic*. RAiSD was run on all brown bear populations to select candidate genes under positive selection exclusive to the ABB. We used the default parameters and computed the *μ statistic* for overlapping sliding windows of 20 SNP with a 1-SNP sliding interval, and then converted these windows into non-overlapping windows of 50 kb using bedtools in order to obtain the same genomic windows as in the other analyses to facilitate comparisons between different results.

We used a second program to identify regions of the genome under selection. In particular, XP-CLR is based on a composite likelihood method and detects selective sweeps in pairs of populations by searching for extreme allele frequency differentiation over genomic windows. Values were calculated per chromosome using xpclr v1.1.2^53^ on the dataset with all samples after quality filtering. We used non-overlapping windows of 50 kb (--size 50000 --step 50000) in the pairwise comparisons.

Finally, we implemented the population branch statistics (PBS) using four populations, as illustrated in Jiang et al.^158^, to look for genomic regions specifically differentiated in the ABB compared to the European and North American samples. We computed the mean pairwise F_ST_ values in non-overlapping 50 kb windows with vcftools, setting to zero the negative indexes and to 0.99 the fixed differences to avoid infinite branch length. Then we applied the Cavalli-Sforza transformation and obtained the PBS values by applying the following equation:

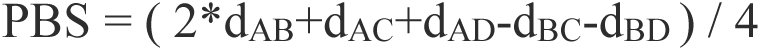

Where *d* is the log-transformed distance between two populations and A,B,C,D are ABB, SBB, Canada and Alaska populations. The windows showing a negative PBS, which can be obtained in case of small sample size, were set to zero as they do not have a biological meaning. This last method was applied to the SV datasets as well. The only procedural difference concerned the F_ST_ calculation that for the SVs was done on each marker rather than in a sliding window fashion.

To identify a potential list of regions under selection and contribute to understanding how selection has affected the genomic variation in ABB, we defined outliers following a similar procedure for each analysis (RAiSD, XP-CLR and PBS) and combined all outlier genes to get a list of candidate genes under selection in the ABB. We first computed the top 1% windows for each analysis. Then, for the SNP dataset we merged outlier windows less distant than 100 kb to extend outlier windows when another outlier window was located in close proximity. Finally, we extracted genes present in these outlier windows or regions by blasting the ABB transcripts to the dog genes. As some analyses were run to get candidate regions under positive selection in the ABB population and others were run as a control in the remaining populations, we then obtained the final list of candidate genes by considering only the outliers in ABB from RAiSD and PBS or in the ABB-SBB comparison from XP-CLR that were not outliers in the other control analyses (i.e., RAiSD for the SBB, Canadian brown bear or Alaskan brown bear, or XP-CLR for the ABB / Canadian brown bear or ABB / Alaskan brown bear). We then performed gene ontology (GO) and human phenotype (HP) enrichment analysis with g:Profiler^159^ for candidate genes identified for all three analyses combined together using the dog gene set. As the GO enrichment analysis might not identify pathways related to selection affecting a few genes of major effect, we also performed an extensive review of the function of the candidate genes present in at least one of the three selection scan analyses.

Similarly, for the PBS analysis on the SV dataset we considered outlier the SVs in the top 1% carrying the derived allele in the ABB population and conducted a separate GO enrichment analysis on g:Profiler using the available polar bear gene set.

We repeated all the selection scans and GO enrichment analysis using the SBB as the target population in order to verify if the signatures of selection we identified in the ABB were the result of a true signal specific to that population. We also computed Tajima’s D in 50 kb windows with vcftools to verify whether the whole ensemble of candidate genes under selection in the ABB were behaving differently from the same number of random genes or intergenic regions while a different pattern could be expected in the SBB.

We decided to focus on one of the phenotypes that distinguishes the ABB from the other brown bear populations, i.e. reduced aggressiveness towards humans. Therefore, we gathered a list of 17 candidate genes under selection (see Results) that we found to have a putative role in the determination of this phenotype, and we analysed them in more detail to possibly unveil the molecular mechanism responsible for the less aggressive attitude of the ABB. After noticing that most of the genetic variation was found in intronic regions rather than in coding portions of the candidate genes (Tab. 1), we applied a series of *in silico* analyses to investigate the potential alteration of splicing at the above-mentioned genes. We started by scanning the genes for SNPs that could alter the acceptor/donor sites recognition during pre-mRNA maturation with SpliceAI ^160^. This program was shown to perform better than many other tools^161^ especially for deep intronic variants because of the large windows provided during the training of its deep learning model^160^) and has already been applied to species other than *Homo sapiens*^162^. We applied the analysis on the maximum window length allowed (i.e. 10 kb) and considered potential splice-altering sites of all the SNPs with delta score > 0.2.

Another feature that can lead to alternative splicing is the alteration of splicing factor site recognition. We used two different tools to explore this process: HOT-SKIP^163^ was developed to score exonic variants for their probability of inducing exon skipping; SpliceAid2^164^ allows to identify RNA-binding proteins target motifs. As these programs are available as web-interface only, we decided to limit the number of SNPs to analyse by considering only the most promising genes and retaining the SNPs with F_ST_ ≥ 0.73, 0.89 and 0.80 between ABB and SBB, Canadian and Alaskan bears, respectively (top 10% in each comparison).

## ACKNOWLEDGEMENTS

This work was supported by the University of Ferrara (Italy) and funded by the MIUR PRIN 2017 grant 201794ZXTL to G.B.; S.T.V. was supported by a Young Researchers (Marie Skłodowska-Curie winners) grant awarded by the Italian Ministry for Universities and Research; P.C. was supported by the European Union—NextGenerationEU National Biodiversity Future Center. We thank the Huck Institutes of Life Sciences and Penn State Altoona for the funding provided for G.F. We also thank Leonardo Gentile and the National Park of Abruzzo Lazio and Molise for their assistance during sampling activities.

## AUTHOR CONTRIBUTION

A.B. and G.B. conceived the study. L.P. and G.B. coordinated and performed sample collection. A.I. performed laboratory work. M.S. and M.Gerdol performed Hi-C data generation and analysis. G.F., R.B., M.Gabrielli, B.S., S.T.V., P.S., L.A., G.P., M.S., A.I. performed the analyses. G.F., R.B., M.Gabrielli, B.S., S.F., S.T.V., P.S., A.I., A.B., G.B. wrote the paper. All authors read, revised, and approved the manuscript.

## DATA AVAILABILITY

The genome raw reads and the genome assembly are available at the National Center for Biotechnology Information (NCBI) with the BioProject no. PRJNA1247790. The genome assembly and annotation are available for download from Zenodo (https://doi.org/10.5281/zenodo.15349716).

