## supplementary figures for "Coexisting with humans: genomic and behavioural consequences in a small and isolated bear population"

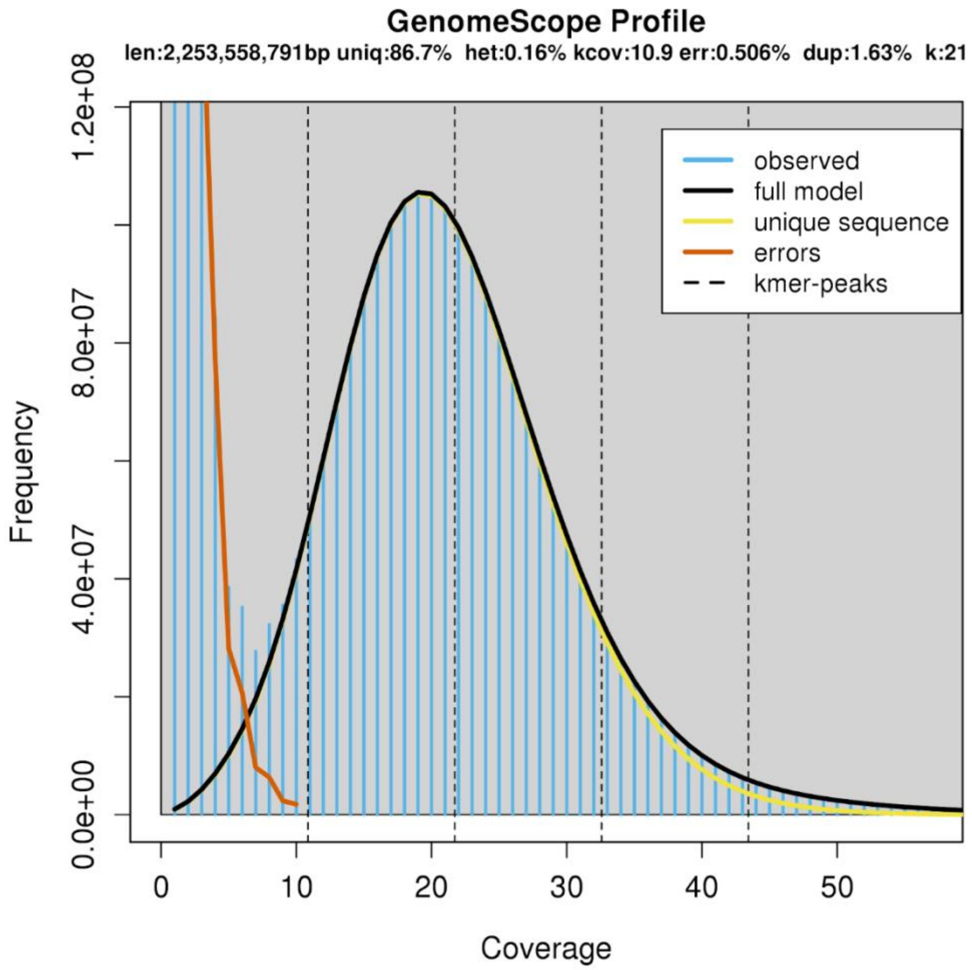


**Supplementary Figure 1:** Genome size estimation from kmers analysis in Genomescope.

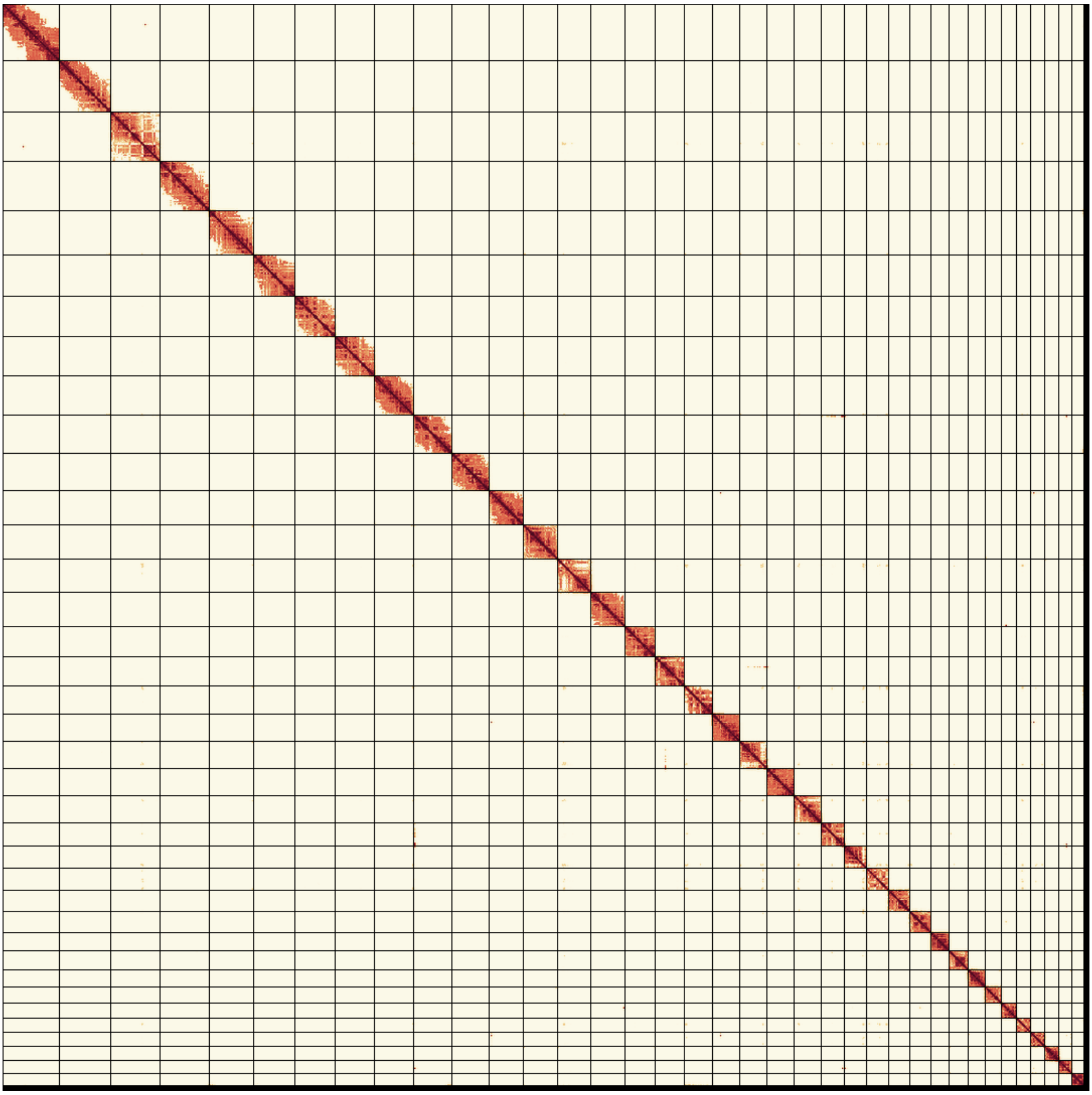

**Supplementary Figure 2:** contact matrix from Hi-C data representing the 36 autosomes and the X chromosome.

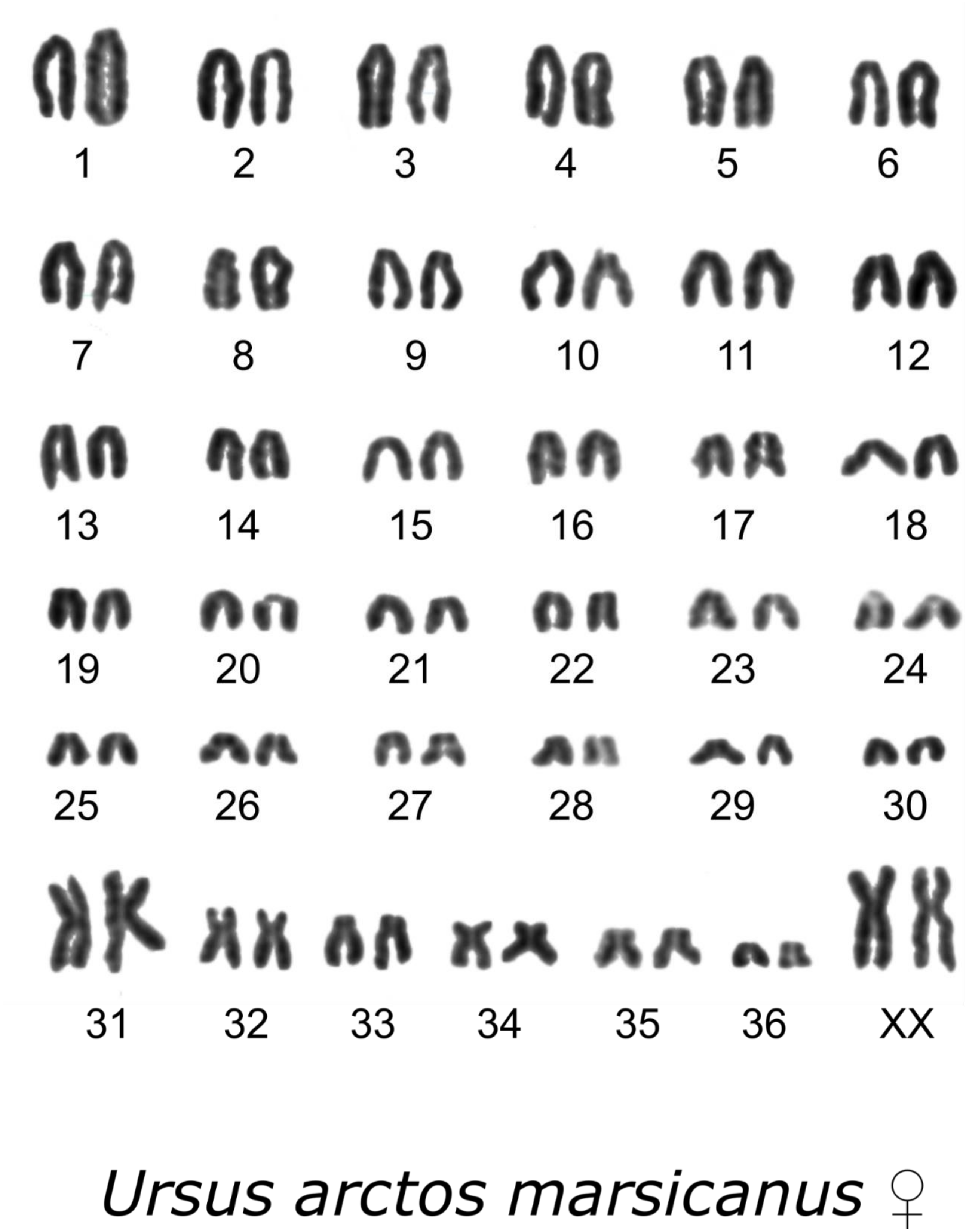


**Supplementary Figure 3:** Apennine brown bear's karyotype including the 36 autosomes and the X chromosome.

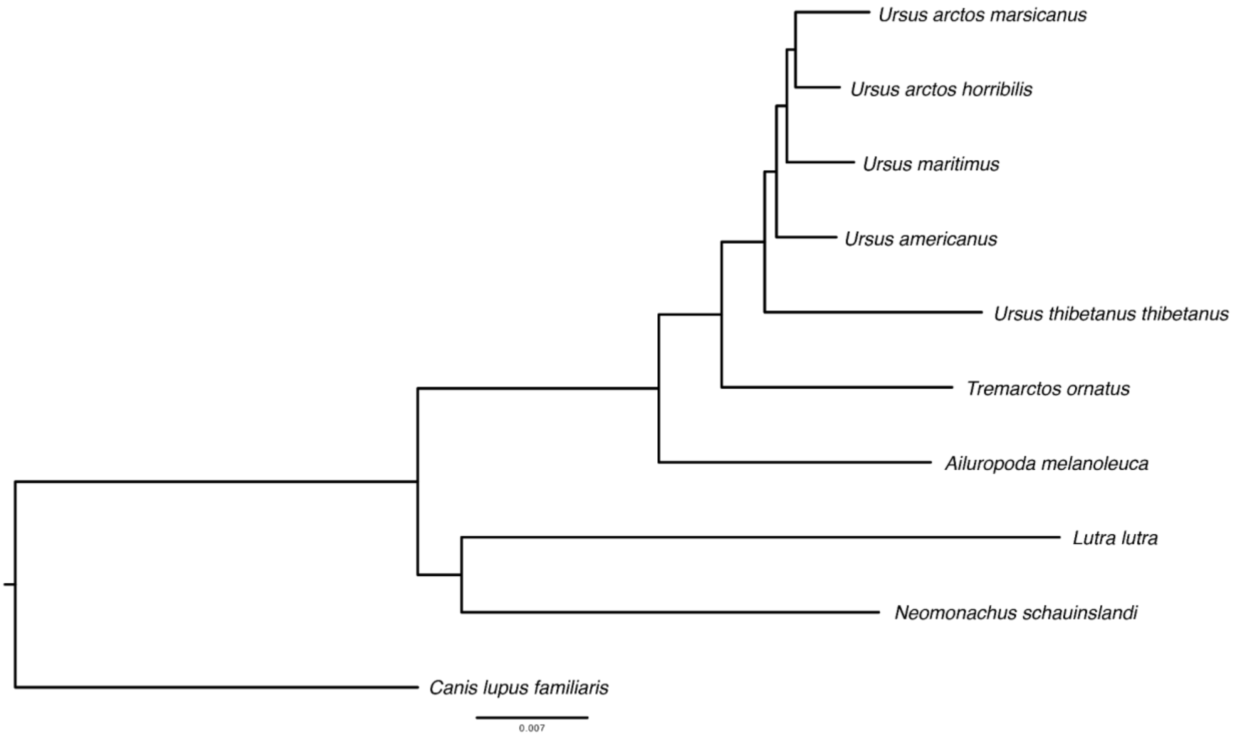


**Supplementary Figure 4:** Phylogenetic tree.

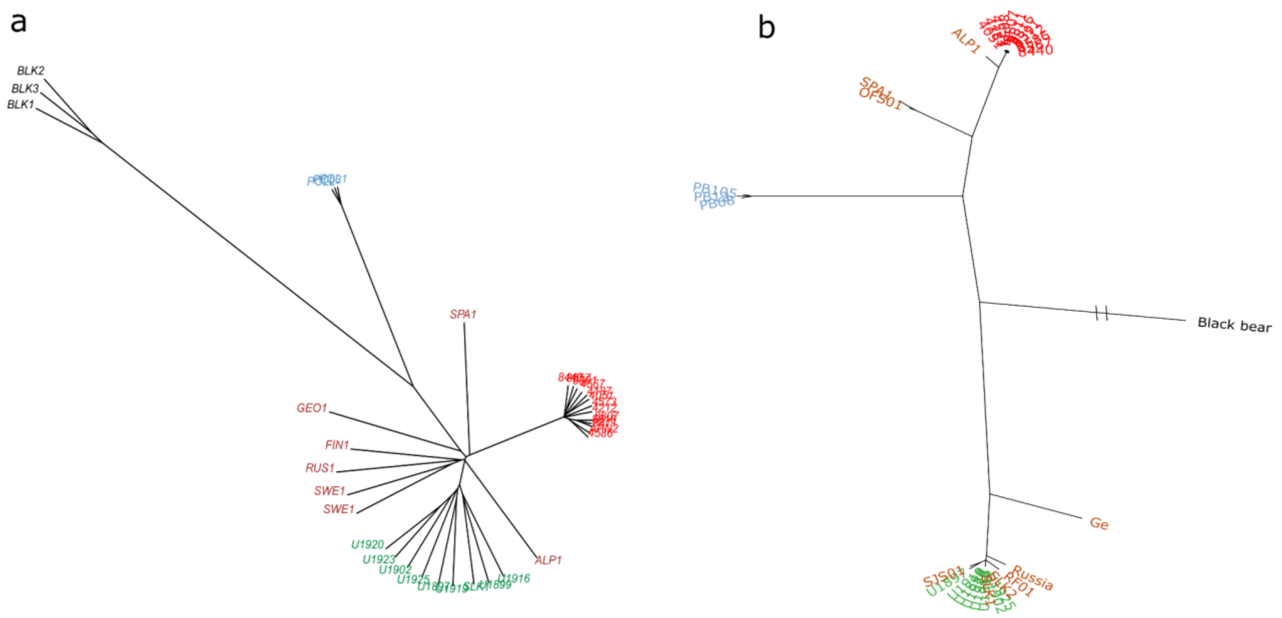


**Supplementary Figure 5:** neighbor joining tree obtained from the analysis of a) nuclear SNPs and b) coding regions and rRNA (12S and 16S). The dataset included several brown bear populations sampled in Eurasia. The Apennine bear (Central Italy) is coloured in red; Slovakia is coloured in green; the rest of the European brown bear samples came from Northern Italy, Spain, Sweden, Finland, Georgia, and Russia and they are all coloured in brown. Polar bears and black bears were also included, coloured in light blue and in black respectively.

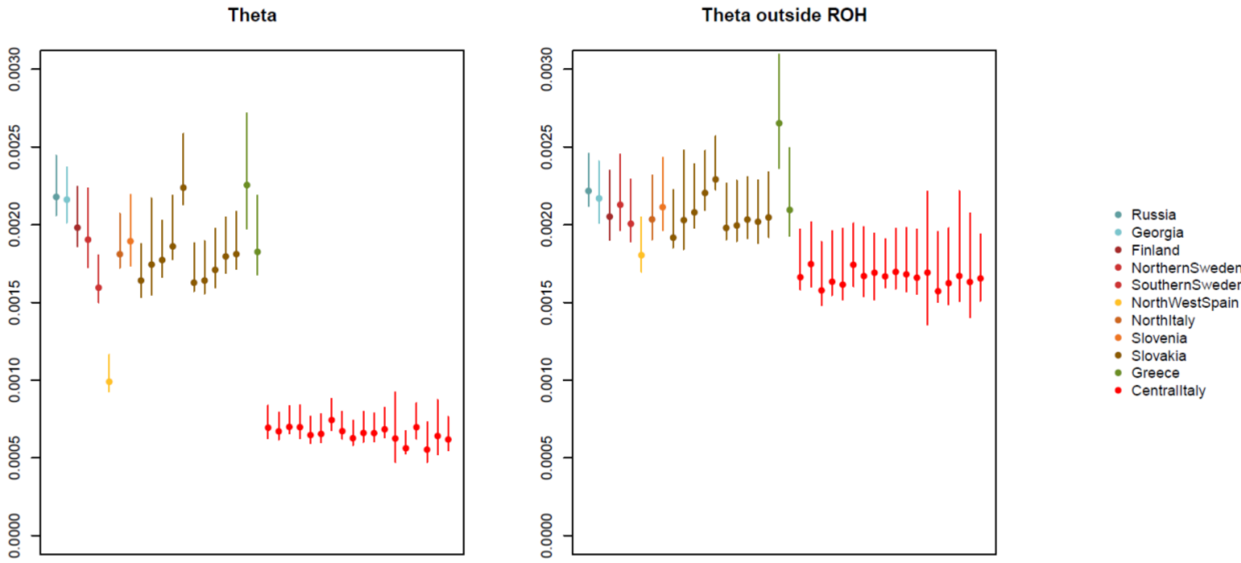


**Supplementary Figure 6:** genetic diversity along the genome of several brown bear populations sampled in Eurasia.

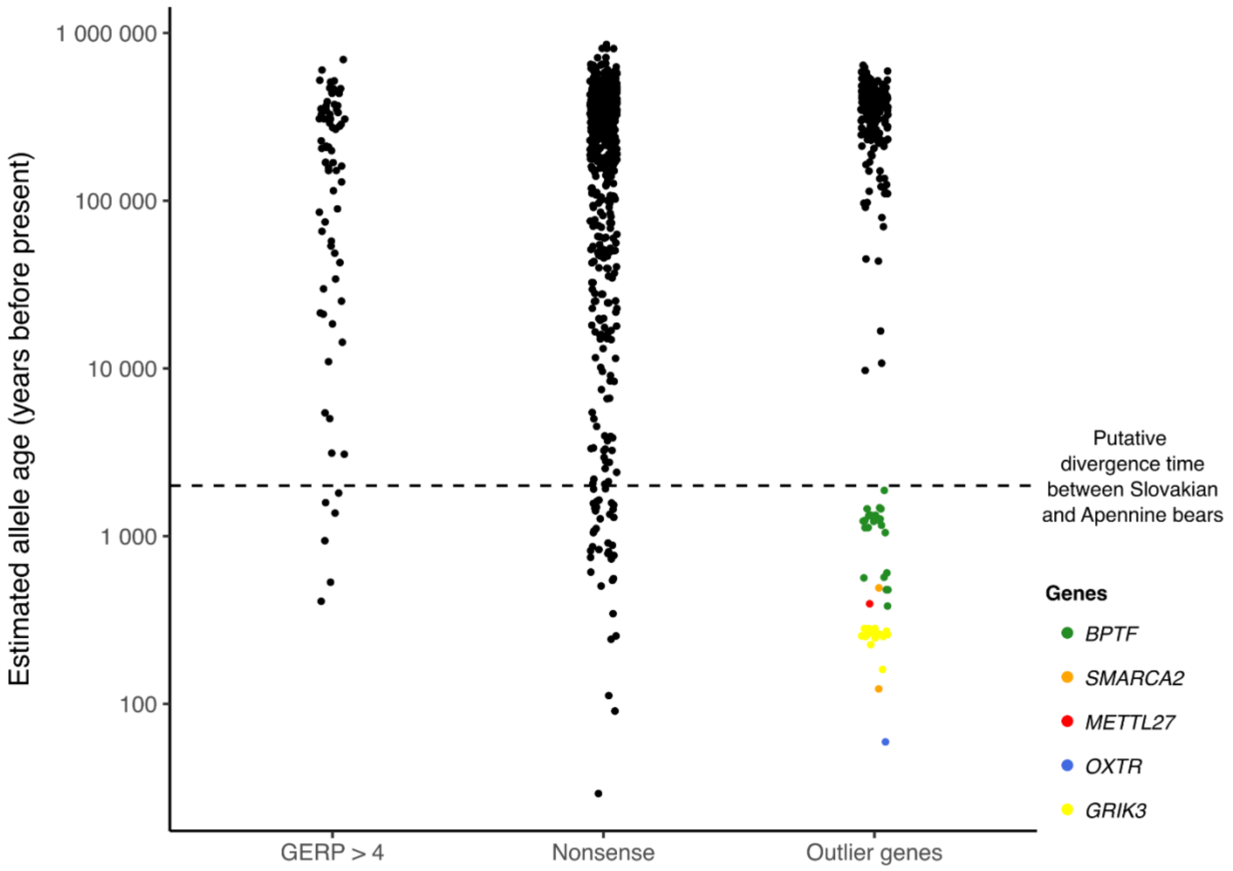


**Supplementary Figure 7:** estimation of allele age for genome-wide highly deleterious variants (GERP > 4), mildly deleterious variants (Nonsense) and for all the segregating sites within the 20 outlier genes under selection with putative influence on behavior. The variants with estimated generation date after the putative divergence time between Slovak and Apennine brown bear are colored according to their corresponding gene.

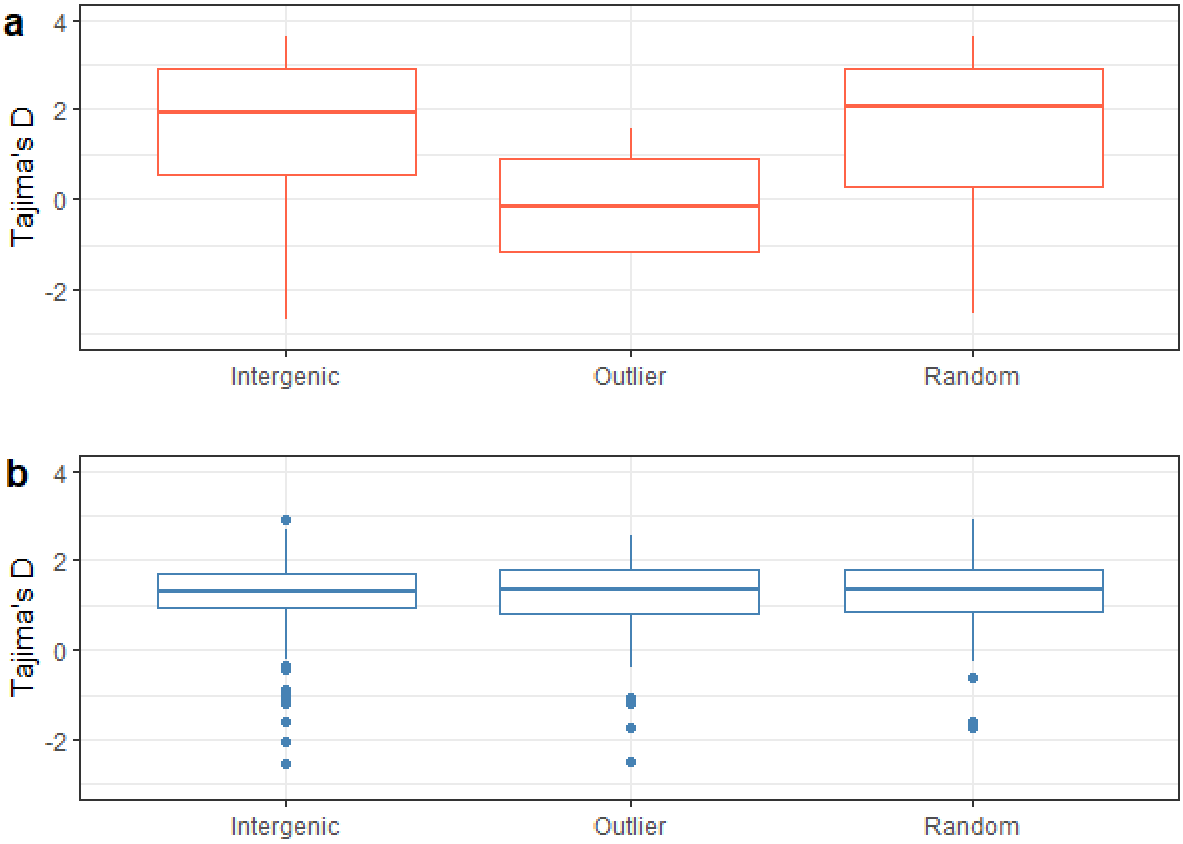


**Supplementary Figure 8:** Tajima's D in Apennine brown bear (a) and Slovak brown bear (b). In the Apennine brown bear the differences were significant for the tests involving the outlier genes compared with the random genes and the intergenic regions (Wilcoxon signed rank test p-values = 5.87e-10 and 1.97e-15 respectively) but were not significant in the comparison between random genes and intergenic regions (Wilcoxon signed rank test p-values = 0.58). No test was significant in the Slovak dataset.
